# The contribution of endocytic mediators to itch transmission

**DOI:** 10.64898/2026.02.13.705766

**Authors:** Marcella de A. Ferreira, Raquel Tonello, Paz Duran, Kai Trevett, Dane D. Jensen

## Abstract

Chronic itch is a major burden, impacting the quality of life for one in four adults, and is closely associated with increased levels of anxiety, depression, and suicide. Despite its widespread burden, the precise mechanisms driving and sustaining chronic itch remain poorly understood. Non-histaminergic itch transmission in the spinal cord relies on the synaptic vesicle (SV) release of gastrin-releasing peptide (GRP) and the subsequent binding to the Gastrin-Releasing Peptide Receptor (GRPR). While SV exocytosis facilitates neurotransmitter release, such as GRP, SV endocytosis, mediated by key proteins including clathrin, adaptor associated kinase 1 (AAK1), and Dynamin (Dnm) are essential for retrieving and recycling SV from the presynaptic membrane to maintain signaling. Building upon evidence that AAK1- and Dnm-mediated endocytosis are viable targets to reverse pain, we characterized the role of these endocytic mediators in the regulation of synaptic transmission in itch pathways. We localized mRNA encoding AAK1, Dnm1 and Dnm3 within GRP positive neurons of the mouse dorsal root ganglia (DRG). Genetic and pharmacological disruption of AAK1 significantly reduced scratching behavior compared to control groups. This anti-pruritic effect correlated with confirmed *Aak1* mRNA knockdown in both the DRG and spinal cord. Similarly, siRNA mediated knockdown of Dnm1 and Dnm3 in the DRGs also reduced scratching behavior. Crucially, these treatments decreased GRP release without altering locomotor activity or anxiety-like behaviors. Together, these findings suggest that the disruption of SV recycling reduces itch related signaling and behavior without affecting normal motor functions, providing a new approach for chronic pruritus.

## Introduction

Pruritus, or itch, is a complex and debilitating sensory modality. The burden of itch is far-reaching, which is associated with increased levels of anxiety, stress, depression, and suicidal ideation. Despite its widespread impact, the precise mechanisms driving pruritus remain poorly understood [19; 27]. While histamine is the main studied mediator of pruritus, current treatment options for chronic itch are limited, particularly for antihistamine-resistant itch, thus, research into non-histaminergic pathways is essential [14]. Itch is defined as an unpleasant sensation that triggers an innate urge to scratch. While initially protective, chronic scratching compromises the skin barrier, releasing inflammatory mediators. If this persists for more than six weeks, it becomes pathological, leading to maladaptive neuroplasticity, including reduced activation thresholds, increased fiber density, and neuroinflammation. These mechanisms mirror those seen in chronic pain, highlighting the shared pathophysiology of itch and pain sensory pathways [12].

In the skin, primary sensory neurons express G protein-coupled receptors (GPCRs) that detect pruritogenic agents at the periphery. The central terminals of these neurons convey pruritogenic signals to the dorsal horn of the spinal cord. The pruriceptor activation on primary sensory neurons results in the release of the neurotransmitter Gastrin-Releasing Peptide (GRP) in the spinal cord. GRP activates its receptor, the Gastrin-Releasing Peptide Receptor (GRPR), which is expressed in excitatory interneurons. Sustained GRP release GRPR signaling is crucial to itch transmission. Prolonged GRP and prolonged activation of GRP-positive neurons evoke depolarization of the interneurons making sustained GRP/GRPR signaling crucial to itch transmission [1; 22].

The sustained release of GRP depends on the synaptic vesicle (SV) cycle [3; 17; 32]. While SV exocytosis releases neurotransmitters into the synapse, SV endocytosis retrieves vesicle components from the presynaptic membrane to maintain a functional SV pool and sustain signaling. Clathrin-mediated endocytosis is the primary mechanism for maintaining the SV pool. Clathrin and adaptor protein 2 (AP2) are the major constitutive proteins of the endocytic SVs. Adaptor-associated kinase 1 (AAK1) recruits clathrin and AP2 to the plasma membrane and phosphorylates the µ2 subunit of AP2 (AP2M1). Finally, dynamin (Dnm), a GTPase, forms a helical collar around the neck of the pit and helps with SV fission. There are three isoforms of Dnm (Dnm1, Dnm2 and Dnm3), with Dnm1 and Dnm3 being highly expressed in the nervous system [11].

The functional importance of these mediators has been established using genetic models. While Dnm3 deletion does not affect synaptic transmission, the combined deletion of Dnm1 and Dnm3 depletes the SV pool and disrupts neurotransmission [24]. Clathrin and Dnm are essential for the endocytosis and endosomal signaling of GPCRs, a mechanism that contributes to neuronal sensitization and nociception [15]. Previous studies have shown that AAK1 and Dnm-mediated endocytosis at nociceptor terminals and inhibition of AAK1 and Dnm reverse pain-like behavior [31]. However, the contribution of these endocytic mediators in itch sensation and signaling has not been characterized. This study investigated the role of endocytic mediators in regulating synaptic transmission within the itch pathways and sought to determine if disrupting SV endocytosis could alleviate itch transmission.

## Methods

### Reagents

All reagents like chloroquine, PBS, DMSO, paraformaldehyde, and sucrose were purchased from Sigma-Aldrich (St. Louis, MO) unless otherwise specified.

### Animals

All experiments were performed in male and female C57BL/6 mice (8-10 weeks, The Jackson Laboratory, Bar Harbor, ME). Following arrival at the animal care facility, mice were allowed to acclimate for at least 1 week and were housed in controlled conditions with a 12-hour light/dark cycle (lights on at 07:00) and a temperature of 22 ± 2°C. Mice had ad libitum access to food and water, with weekly cage changes. Mice were housed in groups of 4–5 and were randomly assigned to experimental groups to avoid cage-related issues. To avoid operator bias, test agents were prepared by a technician, and the investigator was blinded to the groups. The group size was determined by a power analysis based on previous similar studies[31] and designed to include both sexes. All procedures adhered to the *Guide for the Care and Use of Laboratory Animals* (NIH) and were approved by the New York University Institutional Animal Care and Use Committee (PROTO202000064). All efforts were made to minimize animal suffering.

### Intrathecal administration of siRNA to mice

Small interfering RNA (siRNA) plasmids targeting mouse Dnm1 (cat#L-043277-01-0005), mouse Dnm3 (cat#L-059061-01-0005), mouse AAK1 (cat#L-065639-00-0005), or nontargeting control (CTR) siRNA (cat#D-001810-10-05) ON-TARGET plus were used (Dharmacon, Lafayette, CO)(Table S1). For in vivo administration, 1.25 µg of siRNA was complexed with polyethyleneimine-based transfection reagent (in vivo-jetPEI, 201-50 G; Polyplus, Illkirch, France). The siRNA-jetPEI mixture was prepared at an 8:1 nitrogen to phosphate (N:P) ratio, as previously described [6]. The siRNA-jetPEI mixture was administered via intrathecal (i.t.) injection (cervical region C4, 5 µL) to anesthetized mice. Injections were performed 2 days before chloroquine injection (intradermal). To assess the knockdown efficiency, AAK1, Dnm1 and Dnm3 expression in the dorsal root ganglia (DRG) at cervical level was analyzed using RNAScope in situ hybridization immediately after the itch behavioral test (approximately 2 days after siRNA i.t. injections)[31].

### Intrathecal administration of endocytosis inhibitors to mice

Besides siRNA, a pharmacological approach to access the inhibition of endocytic mediators in itch transmission was used. LP935509 (AAK1 inhibitor, cat#HY-117626, MedChemExpress; 10 µg/5 µL), Dyngo4a (Dnm inhibitor, 50 nM), PitStop2 (clathrin inhibitor, 50 nM), (trademarks of Children’s Medical Research Institute, Newcastle Innovation and Freie Universitat Berlin), or vehicle (PBS, 5% DMSO/PBS) was injected i.t. (5 µL, C4) into anesthetized mice. The inhibitors were injected 1 hour before chloroquine injection (intradermally).

### Itch behavioral test

Mice of both sexes were gently anesthetized with isoflurane, and their right napes were shaved. Mice that had received the siRNA previously were given a 10 µL injection of the pruritogen chloroquine (10 mM) into the right nape. Mice that received the pharmacological inhibitors were given the i.t. injection and after 1 hour 10 µL of 10 mM chloroquine in saline was injected intradermally into the right nape of each mouse. Immediately following the chloroquine injection, mice were placed in a Behavioral Spectrometer (Biobserve, Wilhelmstr, Germany). The spectrometer comprised a 40-cm^2^ arena with a CCD camera mounted in the center of the ceiling and a door aperture in the front area of the arena. Movement was assessed by a floor mounted vibration sensor and 32 wall mounted infrared transmitter and receiver pairs. Mice were individually placed in the center of the behavioral spectrometer, and their behavior was recorded for 30 minutes and analyzed using a combination of video tracking analysis (Viewer3, BiObserve, Sankt Augustin, Germany) and vibration analysis [4]. Total distance traveled in the open field (number of visits to a central area), average velocity of locomotion (centimeters/second), track length (centimeter), ambulation (% activity), time engaged in grooming (minutes), and wall distance (centimeter) were recorded and analyzed as described. Scratching behavior was analyzed by the experimenter offline after the record. Each scratching bout was defined as lifting the hind paw up, a rapid brushing of the back nape by the hind paw and returning the hind paw to the floor.

### Collection of mouse tissue

Immediately after itch behavioral test, mice were anesthetized with 5% isoflurane and then transcardially perfused, first with phosphate-buffered saline (PBS), followed by 4% paraformaldehyde in PBS. The cervical DRGs and spinal cord were then carefully dissected. Excised tissues were fixed in 4% paraformaldehyde in PBS at 4°C for 4 hours. Following fixation, both tissues were cryoprotected in PBS + 30% sucrose at 4°C for 24 hours. Subsequently, tissue was embedded in Optimal Cutting Temperature (OCT) compound (Tissue Tek, Torrance, CA). Frozen sections, 10-12 µm thick, were prepared and mounted onto Superfrost Plus slides (Fisher, Suwanne, GA). These sections were then air-dried for 15 minutes and stored at −20°C until further use.

### Quantitative real-time reverse transcription polymerase chain reaction

RNA was isolated from snap frozen mouse tissues using Total RNA Purification Plus kit (cat#48400 Norgen Biotek, Canada). cDNA was prepared using MultiScribe Reverse Transcriptase (cat#4311235 Thermo Fisher). cDNA (50 ng) was amplified for 40 cycles by quantitative real-time reverse transcription polymerase chain reaction (qRT-PCR) with Taqman probes (ThermoFisher) for AAK1 (Mn01183675_m1), Dnm1 (DNM1, Mn00802468_m1), Dnm3 (DNM3, Mn00554098_m1), and GAPDH (Mn99999915_g1), on a Quant-Studio 3 Real-Time PCR System, and TaqMan Fast Advanced PCR Mastermix (cat#4444556 Thermo Fisher) using manufacturers protocol. Samples were run in triplicate and normalized by GAPDH expression. The relative expression ratio per condition was calculated using the 2^ΔΔCT^method [18].

### RNAScope in situ hybridization and immunofluorescence

The RNAScope system (Advanced Cell Diagnostics, Newark, CA) was used by the manufacturer’s directions for fresh-frozen tissue except for the omission of the initial on-slide fixation step. Probe hybridization and detection using the Multiplex Fluorescent Kit v2 followed the manufacturer’s directions. Probes to Mm-Aak1 (#1097711-C1), Mm-Dnm1 (#446931-C3) and Mm-Dnm3 (#451841-C2) (mouse) were used. Sections were incubated with TSA VividTM Fluorophore 570 (1:1500, cat#323271, Advanced Cell Diagnostics) for detection. To detect total neurons, hybridized slides were incubated with NeuroTrace 500/525 Green Fluorescent Nissl Stain (1:500, cat#N21480, Invitrogen, Waltham, MA) (10 minutes, RT). To identify itch-related neurons and total neurons, hybridized slides were incubated overnight at 4°C with primary antibodies targeting rabbit anti-gastrin-releasing peptide (GRP) (1:100, cat#A6380, ABclonal, Wuhan, China) and guinea pig anti-NeuN (1:500), respectively. Slides were washed and incubated with goat anti-rabbit Alexa Fluor 488 (1:1000, ThermoFisher) and with goat anti-guinea pig Alexa Fluor 647 (1:1000, ThermoFisher) (1 hour, RT). Slides were washed and incubated with DAPI, 49,6-diamidino-2-phenylindole (1 mg/mL, 5 minutes) and mounted in ProLong Gold Antifade (cat#P36980, Thermo Fisher). Sections were observed using a Leica SP8 confocal microscope with HCX PL APO 20x (NA=0.75) and 40x (NA=1.3) oil immersion objective.

### RNAScope quantification

AAK1, Dnm1 and Dnm3 were localized by RNAScope. Confocal images were analyzed using Fiji ImageJ (NIH) according to ACD Bio-Techne Technical Note and by the model developed previously [31]. Briefly, neurons were identified using the Nissl or the NeuN channel and a mask was created by applying a threshold (Moments) and selecting neurons 50 µm^2^ or larger. Regions of interest were overlaid on the original micrograph, and the number of dots per area were quantified for AAK1, Dnm1, and Dnm3 respectively. Results are expressed as dots per square millimeter tissue. A total of 3 images (20x objective) were analyzed for each mouse (N = 4 mice/groups; 12 images analyzed per experimental group).

### Gastrin Releasing Peptide (GRP) release

siRNA plasmids targeting mouse Dnm1 and Dnm3 combined, or mouse AAK1, or nontargeting control were administered intrathecally to adult female and male mice. Two days after, mice were anesthetized with 5% isoflurane and then decapitated. Two vertebral incisions were made to expose the spinal cord. Pressure was applied with a PBS-filled syringe inserted into the vertebral foramen to extract the spinal cord. The spinal cord was placed in standard Tyrode solution (119 mM NaCl, 2.5 mM KCl, 2 mM MgCl_2_, 2 mM CaCl_2_, 30 mM glucose, and 25 mM HEPES (pH 7.4) ~310 mOsm) and was left to recover for 5 min at 37° C before starting the treatments. Baseline treatment involved bathing the spinal cord in fresh standard Tyrode solution (700 µL) for 5 minutes at 37° C. Immediately after, the baseline fraction was collected and the spinal cord was treated with an excitatory solution (700 µL), consisting of 90 mM KCl, for 5 minutes at 37° C. These fractions were collected for measurement of GRP release. Samples were immediately stored in a −80° C freezer. The concentration of GRP released into the buffer was measured by enzyme-linked immunosorbent assay using a commercially available ELISA kit (cat# MBS268207; MyBiosource, San Diego, CA), following the manufacturer’s instructions.

### Statistical analysis

The group size for each experiment was based on previous studies [23; 31]. Data are presented as mean ± SEM. Groups of n = 8 mice (4 male and 4 female mice) were studied. Differences were assessed using the Student 2-tailed t test for 2 comparisons and 1-way or 2-way analysis of variance (ANOVA) and Sidak, Tukey, Newman–Keuls or Dunnett post hoc tests for multiple comparisons. P < 0.05 was considered significant at the 95% confidence level. Sample sizes and statistical tests are specified in figure legends.

## Results

### Adaptor-associated kinase 1, Dynamin 1 and Dynamin 3 are expressed in itch-related neurons of dorsal root ganglia and dorsal horn of the spinal cord

AAK1, Dnm1 and Dmn3 are highly expressed in neurons at DRG and spinal cord [31] and our results confirm these data. We used RNAScope in situ hybridization to colocalize endocytic mediators within itch-related neurons in DRG and spinal cord of mice. Itch related neurons were identified by the co-expression of the neuronal marker NeuN and GRP (GRP+ve). *Aak1* mRNA was detected in 90% of GRP+ve neurons of mouse DRG (Figs. 1A and 1D). *Dnm1* mRNA was detected in 80 % of GRP+ve neurons in mouse DRG (Figs. 1B and 1D) and *Dnm3* mRNA was detected in 90% of GRP+ve neurons in mouse DRG (Figs. 1C and 1D). While *Aak1, Dnm1* and *Dnm3* mRNAs were primarily expressed in the deeper dorsal horn neurons of the spinal cord, they were also observed in a subset of neurons whose cell bodies resided in the superficial laminae (LI, LII, and LIII) within GRP+ve neurons (Supplementary Fig. 1). Thus, we confirmed that AAK1, Dnm1 and Dnm3 are expressed in positive neurons for GRP, a subpopulation of neurons related to itch transmission.

**Figure 1.**
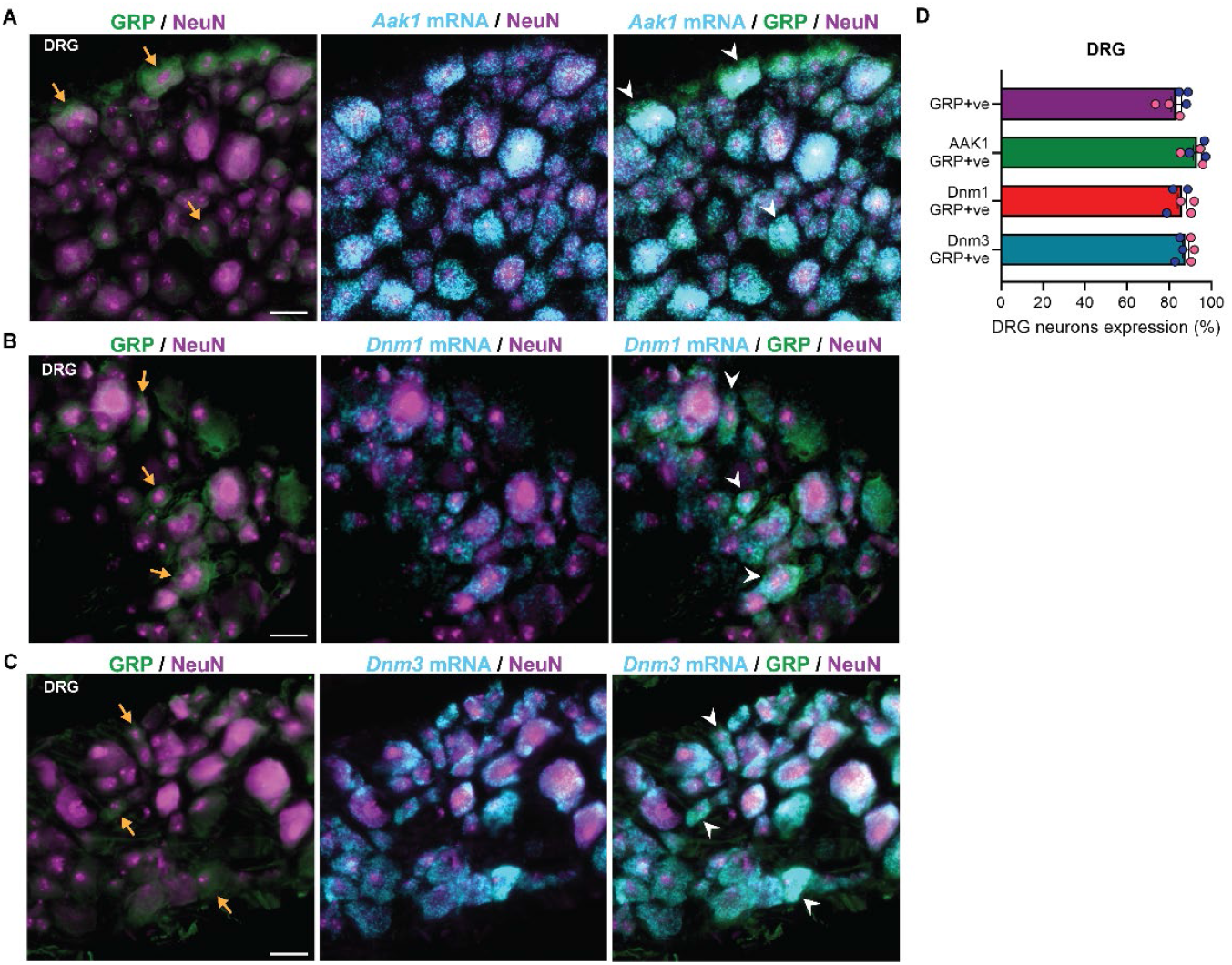
Localization of *Aak1, Dnm1* and *Dnm3* mRNA in itch-related neurons in DRG. Immunofluorescence detection of GRP and NeuN and RNAScope detection of *Aak1* (**A**), *Dnm1* (**B**) and *Dnm3* (**C**) mRNA in mouse DRG. Scale bar 50 µm. (**D**) Percentage of mouse DRG neurons expressing GRP in total neurons, and Aak1, Dnm1 and Dnm3 in GRP+ve neurons (NeuN). Hybridized positive neurons (%). Representative images, n = 6 mice per group (3 male and 3 female). Yellow arrows indicate GRP in total neurons, and white arrows indicate mRNA expression within DRG GRP+ve neurons, blue circles represent male and pink circles represent female mice. Scale bar 50 µm. GRP, gastrin-releasing peptide; DRG, dorsal root ganglia; Aak1, adaptor associated kinase 1; Dnm, dynamin.

### Adaptor-associated kinase 1 siRNA knockdown in dorsal root ganglia and spinal cord reverses itch sensation and scratching

To explore the contribution of AAK1 to itch transmission, AAK1 siRNA or mismatched control (CTR) was administered to mice via cervical i.t. injection. RNAScope analysis subsequently confirmed that AAK1 siRNA successfully reduced the expression of *Aak1* mRNA in DRG neurons by 50 ± 2% after 2 days compared to CTR siRNA (Figs. 2A and 2B). Expression of *Aak1* mRNA in spinal cord neurons also decreased by 72 ± 9% 2 days after AAK1 siRNA compared with CTR siRNA (Figs. 2C and 2D).

**Figure 2.**
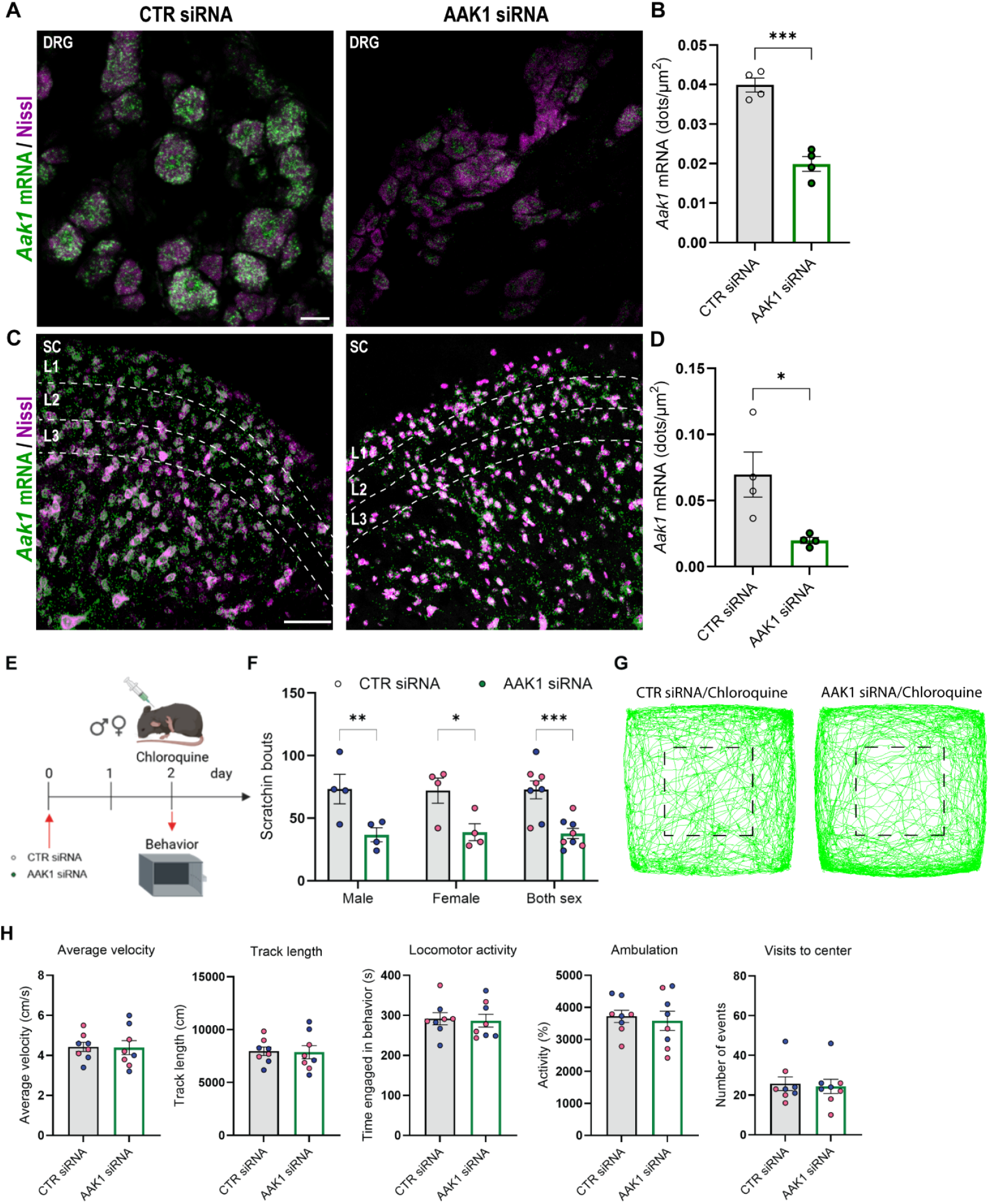
AAK1 siRNA knockdown and scratch behavior on chloroquine-induced itch. **A-D.** RNAScope localization and quantification of *Aak1* mRNA expression in mouse DRG (A, B) and spinal cord (C, D) 2 days after Aak1 or CTR siRNA intrathecal administration. Scale bar 50 μm. n = 4 (B and D). **E**. Experimental behavior timeline. **F**. Scratching induced by chloroquine (10mM/10µL) recorded for 30 minutes 2 days after Aak1 or CTR siRNA. **G**. Representative images of the track recorded are shown with visits to the center area marked by a black square. **H**. Non-evoked behavior in chloroquine-induced scratch recorded for 30 minutes 2 days after Aak1 or CTR siRNA. n = 8 mice per group (4 male and 4 female), blue circles represent male and pink circles represent female mice. Data shown as mean ± SEM. *P<0.05, **P<0.01, ***P<0.001, vs CTR siRNA group, parametric unpaired 2-tailed t test.

Itch behavior was assessed two days following intrathecal AAK1 siRNA injection. To evaluate a histamine-independent pathway with GPCR activation and excitation of sensory neurons, itch was induced by an intradermal injection of chloroquine (10 mM/10 µL) to the nape. After chloroquine administration, mice were immediately placed into a behavioral spectrometer and recorded for 30 minutes where behavior such as scratching bouts, locomotor, exploratory and anxiety-like behaviors were measured (Fig. 2E).

Intradermal chloroquine injection significantly increased the number of scratching bouts, considered as an itch behavior. AAK1 siRNA partially reduces scratching behavior, decreasing it by 48 ± 6%, with both male and female mice showing similar effects compared to CTR siRNA (Fig. 2F).

To evaluate off target effects of siRNA mediated reduction of AAK1, locomotor, exploratory, and anxiety-like behaviors were assessed at the same time as the scratching behavior. AAK1 knockdown by siRNA did not affect average velocity, track length, locomotor activity, ambulation or visits to center when compared with CTR siRNA (Figs. 2G, 2H and Supplementary Fig2A). AAK1 downregulation in DRG and spinal cord neurons effectively suppressed scratching behavior without altering normal behavioral responses or inducing notable side effects.

### Adaptor-associated kinase 1 inhibitor reverse itch sensation and scratching

AAK1 is a validated target for non-opioid pain management as pharmacological inhibition of AAK1 mediated endocytosis effectively disrupts nociceptive transmission [16; 31]. Given the shared neural pathways between pain and pruritus and the effectiveness of siRNA mediate knockdown of AAK1 in inhibiting itch responses, we sought to determine if AAK1 inhibitors could attenuate itch signaling. To test this, we utilized LP935509, a potent and selective AAK1 inhibitor in the chloroquine induced scratching model. LP935509 (10 µg/5 µL) or vehicle (control) was administered (i.t. cervical injection) 1 hour before chloroquine intradermal injection (Fig. 3A). Intradermal injection of chloroquine significantly increased the number of scratching bouts in vehicle-treated mice. The AAK1 inhibitor, LP935509, decreased scratching by 62 ± 4% compared to the control group in both male and female mice. Critically the antipruritic effects of LP935509 occurred without any alteration in average velocity, track length, locomotor activity, ambulation or visits to center behavioral patterns of the mice (Figs. 3B, 3C, 3D and Supplementary Fig2B).

**Figure 3.**
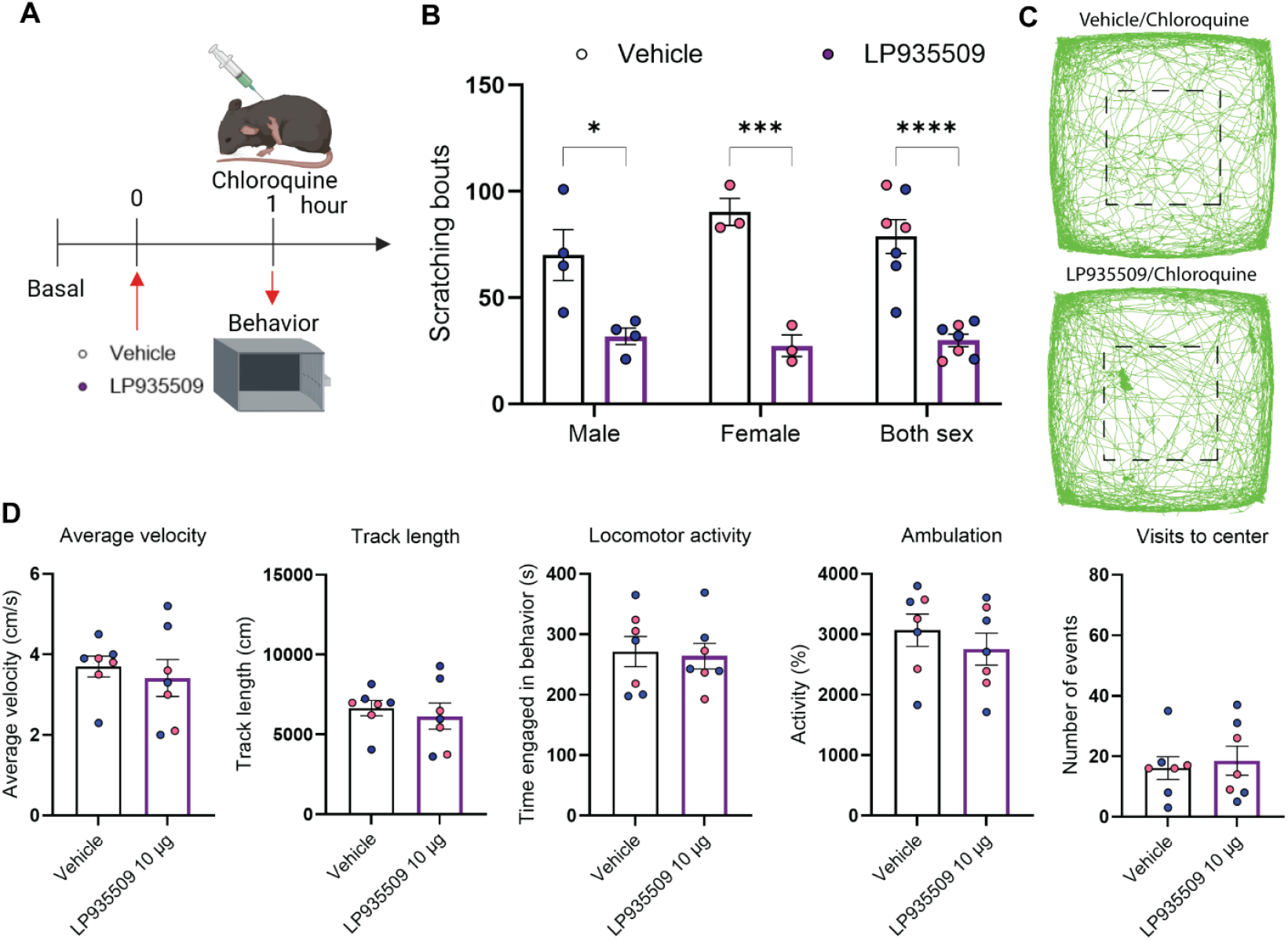
Effect of AAK1 inhibitor on chloroquine-induced itch. **A.** Experimental timeline. **B**. Scratching induced by chloroquine 1 hour after intrathecal administration of AAK1 antagonist (LP935509 10µg/5µL). **C**. Representative images of the track recorded are shown with visits to the center area marked by a black square. **D**. Non-evoked behavior in chloroquine-induced scratch recorded for 30 minutes 1 hour after AAK1 antagonist (LP935509 10 µg, i.t.). n = 7 to 8 mice per group (4 male and 3 female mice), blue circles represent male and pink circles represent female mice. Data shown as mean ± SEM. *P<0.05, ***P<0.001, ****P<0.0001, vs vehicle group, parametric unpaired 2-tailed t test.

### Dynamin siRNA knockdown in dorsal root ganglia reverses itch sensation and scratching

To explore the contribution of Dnm isoforms on itch transmission, mice received a cervical i.t. injection of either Dnm1 siRNA, Dnm3 siRNA, or a mismatched CTR siRNA. RNAScope analysis confirmed the successful reduction of target mRNA expression in DRG neurons 2 days post-injection (Figs. 4A and 4E). Specifically, Dnm1 siRNA reduced *Dnm1* mRNA expression by 57 ± 7%, while Dnm3 siRNA reduced *Dnm3* mRNA expression by 52 ± 6% when compared to the CTR siRNA (Figs. 4B and 4F). While Dnm1 siRNA injection did not affect *Dnm1* mRNA expression in spinal cord neurons (Figs. 4C and 4D), Dnm3 siRNA injection showed a trend towards decreased *Dnm3* mRNA levels in the spinal cord neurons (p=0.0947) (Figs. 4G and 4H).

**Figure 4.**
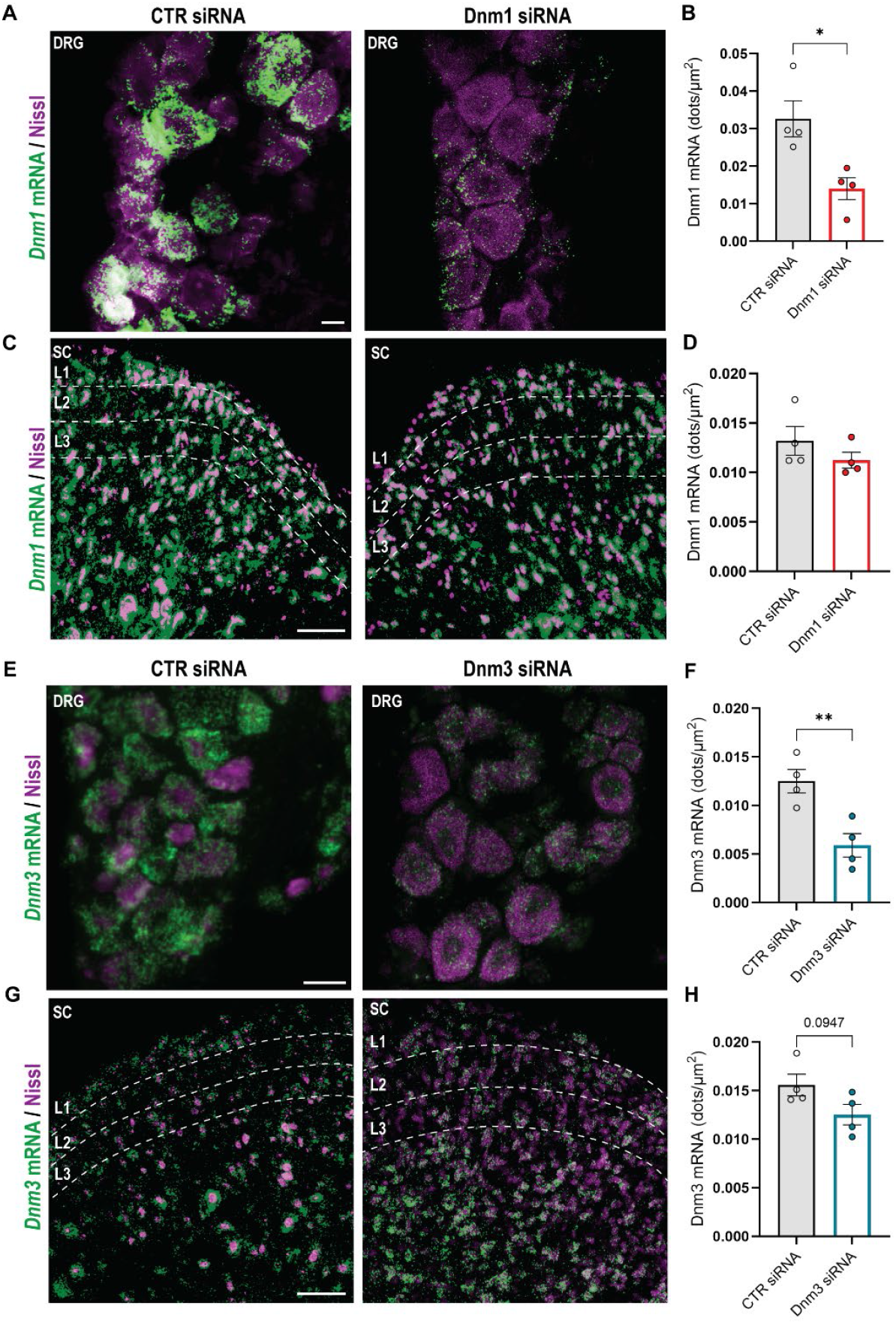
Dnm1 and Dnm3 siRNA knockdown. **A-D.** RNAScope localization and quantification of *Dnm1* mRNA expression in mouse DRG (A, B) and spinal cord (C, D) 2 days after intrathecal administration of Dnm1 or CTR siRNA. **E-H**. RNAScope localization and quantification of *Dnm3* mRNA expression in mouse DRG (E, F) and spinal cord (G, H) 2 days after intrathecal administration of Dnm3 or CTR siRNA. Scale bar 50 μm. n = 4 mice per group. Data shown as mean ± SEM. *P<0.05, vs CTR siRNA, parametric unpaired 2-tailed t test.

Itch behavior was assessed two days after i.t. injections of Dnm1 and Dnm3 siRNA (Fig. 5A). Following chloroquine injection, the number of scratching bouts significantly increased in the CTR siRNA group. In contrast, the Dnm1 siRNA group exhibited a notable reduction in scratching behavior, decreasing by 46 ± 11% when data from both male and female mice were combined. Although, when analyzed separately by sex, just the male Dnm1 siRNA group had a significant decrease in scratching bouts (Fig. 5B). Even more pronounced, Dnm3 siRNA robustly decreased scratching bouts by 69 ± 4%, a significant reduction observed regardless of whether male and female mice were analyzed separately or together (Fig. 5E).

**Figure 5.**
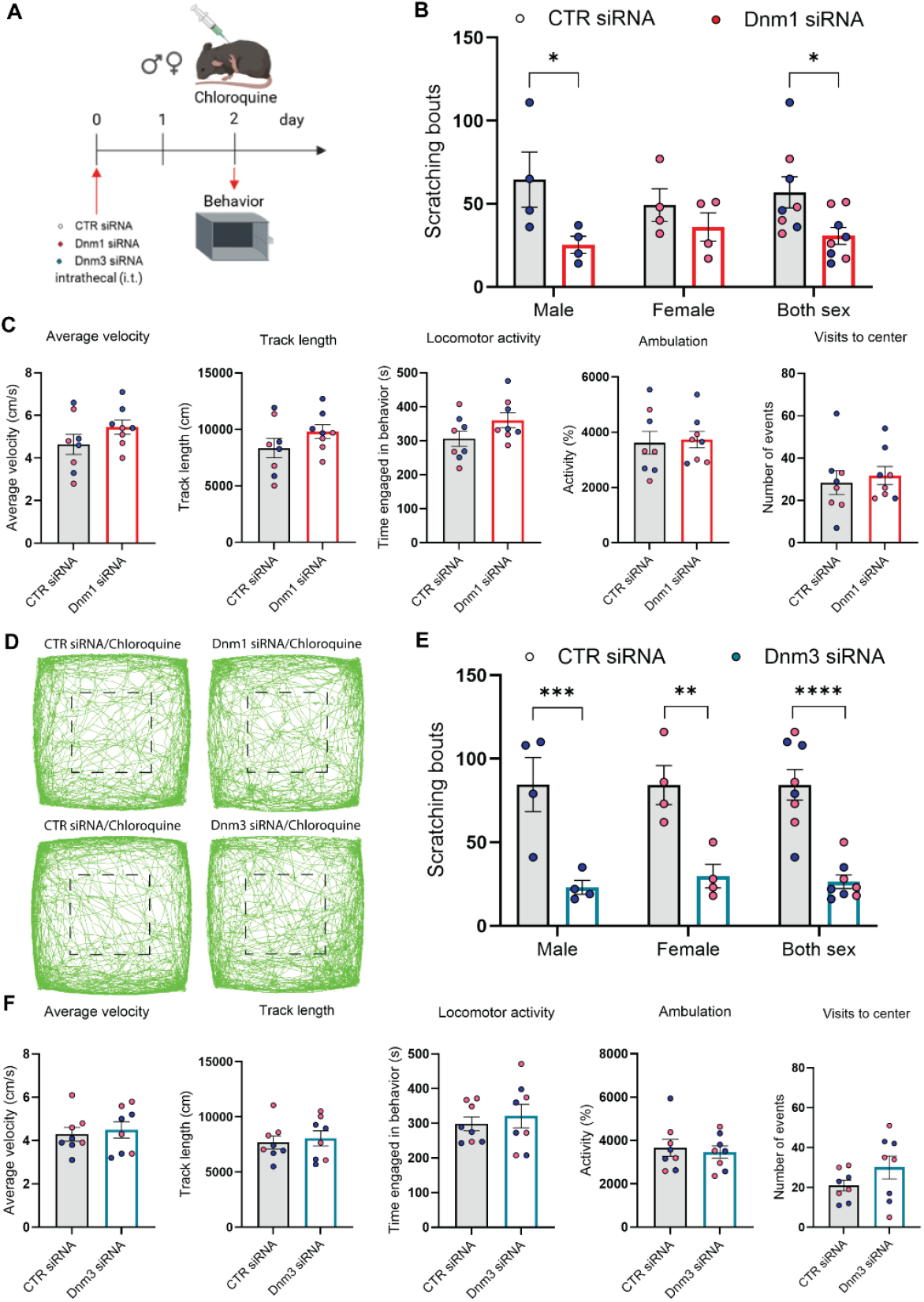
Effect of Dnm1 and Dnm3 siRNA knockdown on chloroquine-induced itch. **A.** Experimental timeline. **B**. Scratching induced by chloroquine 2 days after intrathecal administration of Dnm1 or CTR siRNA. **C**. Non-evoked behavior in chloroquine-induced scratch recorded for 30 minutes at 2 days after Dnm1 or CTR siRNA. **D**. Representative images of the track recorded are shown with visits to the center area marked by a black square. **E**. Scratching induced by chloroquine 2 days after Dnm3 or CTR siRNA. **F**. Non-evoked behavior in chloroquine-induced scratch recorded for 30 minutes at 2 days after Dnm3 or CTR siRNA. n = 8 mice per group (4 male and 4 female mice), blue circles represent male and pink circles represent female mice. Data shown as mean ± SEM. *P<0.05, ***P<0.001, ****P<0.0001, vs CTR siRNA group, parametric unpaired 2-tailed t test.

Again, to assess off target effects of Dnm1 or Dnm3 siRNA mediated knockdown, locomotor, exploratory, and anxiety-like behaviors were evaluated using a behavioral spectrometer immediately following chloroquine injection. Critically, neither Dnm1 nor Dnm3 siRNA significantly affected average velocity, track length, locomotor activity, ambulation or visits to the center when compared to the control (CTR) siRNA groups (Figs. 5C, 5D, 5F, Supplementary Fig2C and Supplementary Fig2D). These findings indicate that Dnm1 and Dnm3 downregulation in DRG neurons effectively suppressed scratching behavior without impairing normal motor functions or behaviors.

### Adaptor-associated kinase 1, Dynamin 1 and Dynamin 3 siRNAs reduces the quantity of GRP release

Finally, we tested whether spinal pruriceptive neurotransmission was decreased by AAK1 and Dnm knockdown. Two days after administering AAK1, Dnm1+Dnm3, or control (CTR) siRNAs to mice via i.t injection, we measured the release of Gastrin Releasing Peptide (GRP), the excitatory neurotransmitter modulating itch transmission, in the spinal cord. We used an *ex vivo* assay where mouse spinal cord is stimulated with a high concentration of KCl to induce membrane depolarization. Baseline and excitatory fractions were collected, and evoked immunoreactive GRP (iGRP) content was measured by enzyme-linked immunosorbent assay (ELISA). Both AAK1 and Dnm1+Dnm3 knockdown significantly reduced the release of evoked iGRP compared to the control condition (Fig. 6; KCl). Furthermore, analysis of baseline release showed a downward trend in both knockdown groups compared to control condition (Fig. 6; baseline). These results demonstrate that knockdown of the endocytic mediators AAK1, or Dnm1+Dnm3 reduces spinal GRP release.

**Figure 6.**
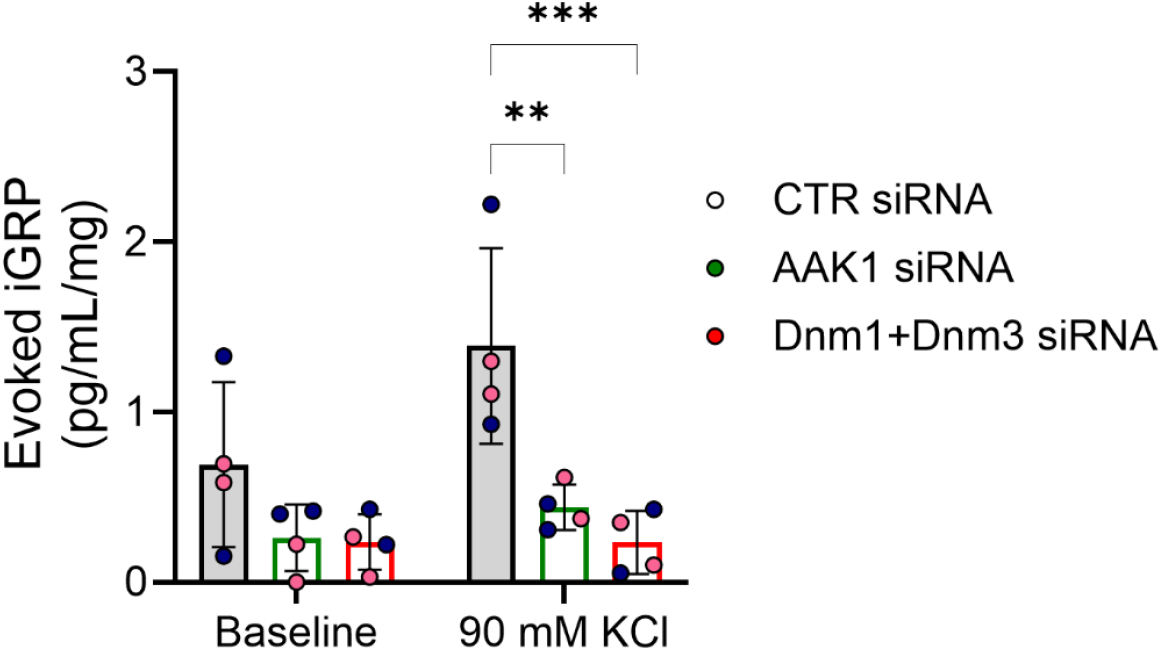
Effect of AAK1, and Dnm1+Dnm3 siRNA knockdown on spinal cord neurotransmitter release. KCl depolarization-evoked GRP release was measured from spinal cords isolated from female and male mice. Bar graphs show normalized immunoreactive GRP levels. n=4 mice per group (4 male and 4 female mice), blue circles represent male and pink circles represent female mice. Data shown as mean ± SEM. **P<0.01, ***P<0.001, two-way ANOVA with a Tukey’s multiple comparisons test.

## Discussion

Previous studies identified AAK1, Dnm1 and Dnm3 as SV recycling mediators in nociceptive pathways and showed that SV recycling is critical for sustained synaptic transmission in pain transmission [16; 31]. As pruritus and nociception use shared neural pathways, we investigated the role of these synaptic mediators, AAK1, Dnm1 and Dnm3, in synaptic transmission in itch pathways. Histamine remains the most extensively studied mediator of itch, however, many cases of chronic itch are antihistamine-resistant, making current treatment options limited [14]. In this study, we investigated itch transmission by using chloroquine to induce a non-histaminergic itch model. While the expression of AAK1 and Dynamins has been documented in human and mouse sensory neurons, we demonstrate here that AAK1, Dnm1, and Dnm3 are highly localized within GRP positive neurons in the DRG and spinal cord, and that they regulate GRP release, suggesting a key role in the itch sensory pathway.

Pruritogens are detected by different GPCRs expressed on primary sensory neurons in the periphery. Pruritogenic GPCRs, on primary sensory neurons, generate signals that activate transient receptor potential (TRP) family of ion channels. These actions initiate neuronal excitation, leading to the central transmission of the itch signal. Neurotransmitters, such as GRP, are released by pruriceptive sensory neuron [2] and by the excitatory spinal interneuron [28]. Sustained GRP release activates the GRPR receptor in the superficial laminae I in the dorsal horn of spinal cord. The GRPR serves as a final common pathway for non-histaminergic itch pathways, as evidenced by previous studies showing that intrathecal administration of GRPR antagonists effectively blocks scratching behavior in mice and the ablation of GRPR positive interneurons regulates scratching behavior in mice [30]. The prolonged release of GRP results in the increased trafficking of GRPR which causes activation of subcellular compartments in the cytosol and nucleus, contributing to sustained signals in the spinal interneurons that underline persistent excitation and chronic itch. Our findings suggest that disrupting SV endocytosis by targeting AAK1 or Dnm1/3 interrupts this signaling cascade by blocking the sustained release of GRP that necessary to open the spinal gate for itch[22].

Specifically, AAK1 facilitates the clathrin-mediated endocytosis of SVs by phosphorylating the µ2 subunit of AP2 [20]. The maintenance of the SV cycle is crucial to sustain neurotransmission of itch sensation as siRNA knockdown of AAK1 in DRG and spinal cord neurons reduced GRP release and blocked scratching behavior in mice. These results were corroborated by the small-molecule AAK1 inhibitor LP935509. Importantly, neither intervention altered basal locomotion or anxiety-like behaviors, supporting the hypothesis that AAK1 is a viable therapeutic target that modulates pathological signaling without disrupting basal neurotransmission [16; 31].

Like AAK1, dynamin is a key component in the SV cycle. Dynamin is a multidomain GTPase which regulates the clathrin-mediated endocytosis by catalyzing the SV fission [20]. siRNA mediated knockdown of Dnm1/3 blocked chloroquine induced scratching in mice. Again, as was shown with AAK1 knockdown, Dnm1/3 knockdown did not interfere in normal non-pruritic behaviors. Dnm1/3 knockdown did not alter locomotor activity or anxiety-like behaviors. These results are in accordance with the literature that shows the maximal Dnm knockdown in mice using a cocktail of siRNAs targeting Dnm1, 2, and 3 (Dnm1+2+3 siRNA), which significantly reduced GRP release, inhibited scratching behavior in both male and female mice without changing locomotor, exploratory nor anxiety-like behaviors. Conversely, intrathecal injection of Dynamin inhibitor, Dyngo4a, led to a significant reduction in scratching bouts similarly in both male and female mice [26].

A key finding of this study is that AAK1 and Dynamin inhibition selectively impacts evoked itch responses while sparing non-evoked behaviors. This suggests an activity-dependent requirement for rapid SV recycling. Only when sensory neurons enter a state of heightened activity, such as during chronic pruritus which requires sustained GRP release, does the system become reliant on the high-efficiency recycling mediated by AAK1 and Dynamin. These results are in accordance with previous nociceptive studies results that showed AAK1 and Dnm are needed for prolonged nociceptive signaling [13; 25; 31].

While global targeting of endocytosis poses risks due to its ubiquitous cellular functions, our results demonstrate that localized disruption (either via siRNA or specific inhibitors) effectively suppresses pathological itch without visible side effects. This distinction is critical, as total Dnm1/3 knockout is lethal [24]. A partial disruption of SV endocytosis mediators appears to be crucial for maintaining basal synaptic transmission while effectively suppressing the excessive synaptic activity involved in itch transmission. Furthermore, while AAK1 knockdown was observed in both the DRG and spinal cord, Dnm1/3 knockdown was largely restricted to the DRG. This localized effect was nonetheless sufficient to block scratching, highlighting the importance of peripheral SV cycling in itch. The lack of spinal Dnm knockdown may stem from limited tissue penetration or the need to target all three isoforms simultaneously. Notably, the sensitivity of RNAscope proved superior to qPCR in detecting these changes, as it allowed for the specific analysis of mRNA within targeted neuronal populations (Suppl. Fig. 3).

The clinical potential of this approach is underscored by existing pharmacology; for example, the AAK1 antagonist LX9211 was safe and well-tolerated in healthy human subjects during Phase 1 clinical trials [5]. Furthermore, the antitumor and antipsychotic drug prochlorperazine enhances cancer immunotherapy by inhibiting Dnm [7; 9]. Recently, other synaptic mediators, such as Endophilin A (EndoA1) and Synaptojanin 1 (Synj1), were investigated in nociceptive transmission context. Endophilin A participates in endocytic membrane retrieval and uncoating in neurons, while Synaptojanin 1 is needed for the neck formation of endocytic pits [8; 21]. Silencing EndoA1 and Synj1 reversed both mechanical allodynia and thermal hyperalgesia in preclinical models of postoperative and cancer pain [23]. Further studies of other synaptic mediators, including EndoA1 and Synj1, are necessary to investigate the participation of these mediators to itch transmission.

Furthermore, existing evidence shows sexual dimorphism in itch transmission. Finally, we addressed potential sexual dimorphism in itch transmission. Despite reports of higher itch prevalence and intensity in females [10; 29] we found no significant sex-based differences in the expression or function of AAK1 and Dynamins in this context. While a non-significant trend was noted in female Dnm1 siRNA groups, this likely reflects experimental variation rather than a divergent biological mechanism. Collectively, our data establish AAK1 and Dynamin isoforms as potent, sex-independent targets for managing chronic pruritus. By disrupting the SV recycling necessary for sustained GRP signaling, these mediators offer a novel strategy to control the sustained release of GRP or itch related neurotransmitter that are released in hyperactive itch pathways without compromising normal motor function.

These findings suggest that disruption of SV recycling reduces itch behavior without compromising normal motor functions. This work establishes endocytic mediators, particularly AAK1 and Dynamin isoforms, as promising novel targets for the management of itch.

## Acknowledgments

We thank Tracy Chiu and Evan Chen for technical assistance. This research was funded by NIH R01NS125413 (DDJ).

## Data Availability

The data that support the findings of this study are available from the corresponding author upon reasonable request.

