## Supplementary material for "The contribution of endocytic mediators to itch transmission": Supp Figures

**Supplemental Figures:**

**Fig S1.**

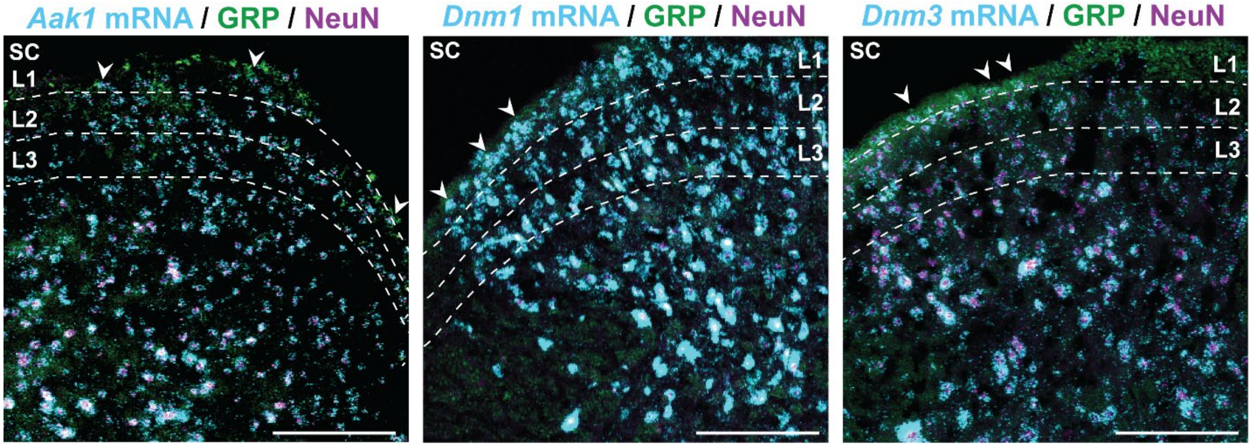


**Fig S1.** Localization of *Aak1*, *Dnm1* and *Dnm3* mRNA in itch-related neurons in the dorsal horn of the spinal cord. Immunofluorescence detection of GRP and NeuN and RNAScope detection of *Aak1*, *Dnm1* and *Dnm3* mRNA in mouse spinal cord. Scale bar 50 µm. Arrows indicate mRNA expression within GRP neurons. Representative images. GRP, gastrin-releasing peptide; Aak1, Adaptor-associated kinase 1; Dnm, Dynamin.

**Fig S2.**

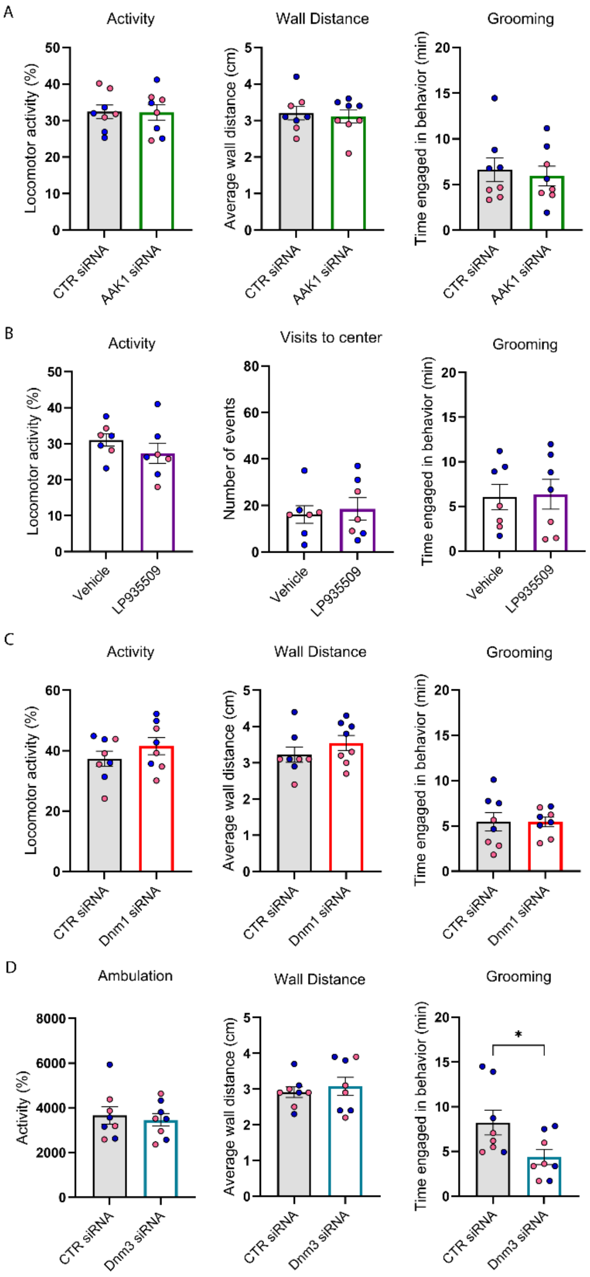


**Fig S2**. Non-evoked behaviors after treatment. Non-evoked behavior in chloroquine-induced scratch recorded for 30 minutes at 2 days after intrathecal administration of AAK1 siRNA (**A**), LP965509 (10µg/5µL) (**B**), Dnm1 siRNA (**C**), Dnm3 siRNA or their respective controls (**D**). n = 8 mice per group (4 male and 4 female mice), blue circles represent male and pink circles represent female mice. Data shown as mean ± SEM. *P<0.05, vs CTR siRNA group, parametric unpaired 2-tailed t test.

**Fig S3**

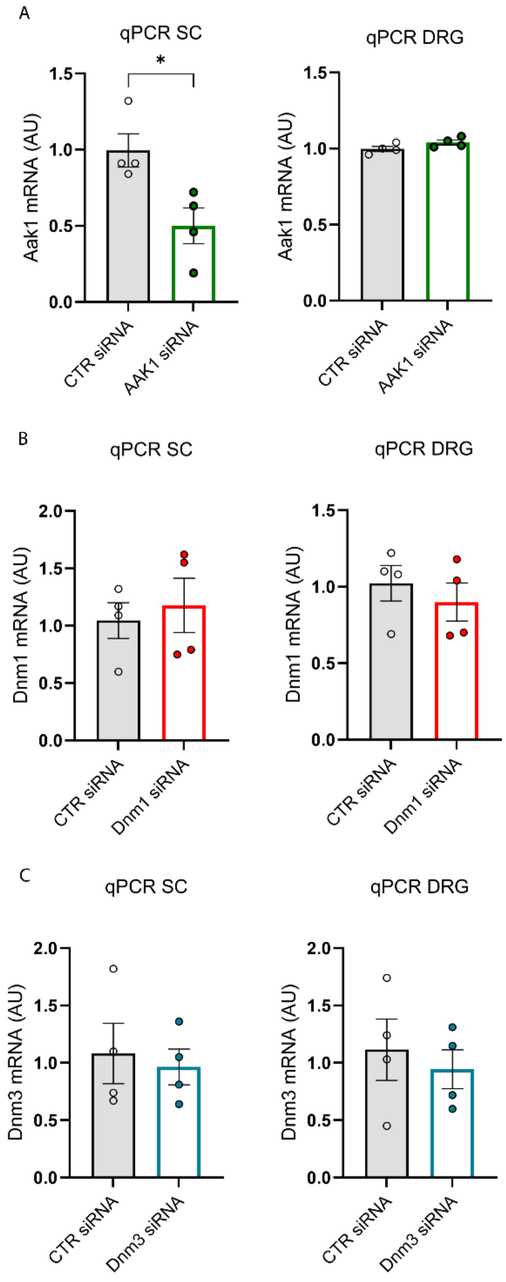


**Fig S3.** Endocytic mediator's mRNA levels after siRNA injections. Expression levels of *Aak1* (**A**), *Dnm1* (**B**), and *Dnm3* (**C**) mRNA in the spinal cord (SC) and DRG of treated mice, determined by qRT-PCR, n=4 mice per group.

**Table S1.** siRNA sequences used to knockdown AAK1, Dnm1 and Dnm3. m, mouse.

| Target | Dharnmacon Sequence |
|---|---|
| *mAAK1* | GAAGGUGGAUUCGCUCUUG,<br>GGACUCAAAUCUCCUGACA,<br>GCAGAUAUUUGGGCUCUAG,<br>AAAUGUGCCUUGAAACGUA. |
| *mDnm1* | GCGUGUACCCUGAGCGUGU,<br>UGGUAUUGCUCCUGCGACA,<br>GGGAGGAGAUGGAGCGAAU,<br>GCUGAGACCGAUCGAGUCA. |
| *mDnm3* | CAACGAAGGCUGACGAUAA,<br>GCUCAGAGUUCCUGCGAAA,<br>GUGAAUGGAACUCGUAUAA,<br>GCAGAAACAGACCGCGUAA |
| *Control* | UGGUUUACAUGUCGACUAA,<br>UGGUUUACAUGUUGUGUGA,<br>UGGUUUACAUGUUUUCUGA,<br>UGGUUUACAUGUUUUCCUA. |
